# MSIsensor-RNA: Microsatellite Instability Detection for Bulk and Single-cell Gene Expression Data

**DOI:** 10.1101/2023.10.09.561613

**Authors:** Peng Jia, Xuanhao Yang, Xiaofei Yang, Tingjie Wang, Yu Xu, Kai Ye

**Affiliations:** MOE Key Lab for Intelligent Networks & Networks Security, Faculty of Electronic and Information Engineering, Xi’an Jiaotong University, Xi’an 710049, China; School of Automation Science and Engineering, Faculty of Electronic and Information Engineering, Xi’an Jiaotong University, Xi’an 710049, China; School of Computer Science and Technology, Faculty of Electronic and Information Engineering, Xi’an Jiaotong University, Xi’an 710049, China; School of Life Science and Technology, Xi’an Jiaotong University, Xi’an 710049, China; Genome Institute, the First Affiliated Hospital of Xi’an Jiaotong University, Xi’an 710061, China

**Author notes:** Corresponding author. (Ye K). Equal contribution (Peng Jia, Xuanhao Yang).

**Keywords:** Microsatellite instability, Gene expression, Single-cell RNA-seq, RNA-seq, Microarray

## Abstract

Microsatellite instability (MSI) is an indispensable biomarker in cancer immunotherapy. Currently, MSI scoring methods by high-throughput omics methods have gained popularity and demonstrated better performance than the gold standard method for MSI detection. However, the MSI detection method on expression data, especially single-cell expression data, is still lacking, limiting the scope of clinical application and prohibiting the investigation of MSI at a single-cell level. Herein, we developed MSIsensor-RNA, an accurate, robust, adaptable, and standalone software to detect MSI status based on expression values of MSI-associated genes. We demonstrated the favorable performance and promise of MSIsensor-RNA in both bulk and single-cell gene expression data in multiplatform technologies including RNA-seq, microarray, and single-cell RNA-seq. MSIsensor-RNA is a versatile, efficient, and robust method for MSI status detection from both bulk and single-cell gene expression data in clinical studies and applications. MSIsensor-RNA is available at https://github.com/xjtu-omics/msisensor-rna.

## Introduction

Microsatellite instability (MSI) refers to hypermutations of microsatellite sites due to inactivating alterations of mismatch repair (MMR) genes in malignancies [1,2]. Currently, MSI is an indispensable pan-cancer biomarker in cancer therapy and prognosis, and it is routinely examined in multiple cancer types, particularly in colorectal cancer (CRC), stomach adenocarcinoma (STAD), and uterine corpus endometrial carcinoma (UCEC) [2□5]. For example, MSI-positive patients are frequently resistant to 5-fluorouracil treatment but exhibit better responses to immune checkpoint blockade treatment [4,5].

In clinical settings, MSI detection mainly relies on the gold standard experimental method, MSI-PCR [6], which is laborious and time-consuming. With the advancement of next-generation sequencing technology, numerous features of genomics, epigenomics, transcriptomics, and histology are investigated, and novel MSI computational algorithms have been developed for a variety of scenarios. [7□21]. Genomics-based methods quantify MSI according to genetic mutations at microsatellite sites, which achieve high accuracy and are becoming popular in clinical MSI detection. For example, MSIsensor [8] achieves a high concordance of 99.4% in detecting MSI on the Memorial Sloan Kettering-Integrated Mutation Profiling of Actionable Cancer Targets (MSK-IMPACT) panel [16]. Epigenomics-based method, MIRMMR, detects MSI using methylation levels in the MMR pathway with the area under the receiver operating characteristic curve (AUC) value of 0.97 [17]. In addition, transcription levels of MSI-associated genes exhibit correlations with MSI, hinting at the possibility of MSI detection using transcriptomics data [15,18,19]. Besides these high-throughput technologies, deep learning algorithms are also applied to hematoxylin and eosin-stained slides to detect MSI [20,21]. However, all these MSI methods detect MSI at a sample level, lacking cell-level measurement of MSI. Recently, single-cell RNA-seq (scRNA-seq) technology has enabled the investigation of cell-specific transcriptomes and shed light on tumor heterogeneity and tumor stages. In particular, the single-cell and spatial transcriptomes enable the dynamic analysis of MSI in the complex tumor microenvironment, such as in metastatic and recurrent cancer [22]. However, current MSI detection methods designed for bulk gene expression data do not perform well on scRNA-seq samples. For example, the only software for gene expression data, PreMSIm [18], provides fixed signatures and a fixed model for all cancers, which limits the wide application of the methods. Moreover, the normalization method in PreMSIm also leads to poor performance with abnormal samples. Here, we have developed MSIsensor-RNA, a robust method for MSI-associated gene detection and MSI evaluation for both bulk gene expression data and single-cell RNA-seq data.

## Method

### Dataset

We downloaded RNA-seq data of 1428 TCGA samples across CRC, STAD, and UCEC from TCGA Research Network (https://portal.gdc.cancer.gov) and obtained their MSI status as determined by gold standards (Table S1). We also obtained 141 RNA-seq samples from the ICGC data portal (https://dcc.icgc.org), and their MSI status was reported by MIMcall [23]. Another 106 RNA-seq samples with their matched MSI status were downloaded from the public publication of the Clinical Proteomic Tumor Analysis Consortium (CPTAC) [24]. We also downloaded microarray data and their MSI status of 1468 samples across CRC and STAD from the GEO dataset (https://www.ncbi.nlm.nih.gov/geo). For scRNA-seq data, we got the gene expression data and their MSI status from 133 CRC samples in two recent publications [25,26].

### Overall design

The pipeline of MSIsensor-RNA consists of data preprocessing, informative gene selection, model training, and model testing (**Figure 1** and Figure S1). First, we preprocessed the expression values of samples from microarray, bulk RNA-seq, and scRNA-seq. Next, we selected an informative gene set for MSI detection from 1428 TCGA samples. Then, these TCGA samples were applied to train a machine-learning model for MSI scoring. Finally, we applied the trained model to independent databases to test the performance of the MSIsensor-RNA for each cancer type.

**Figure 1.**
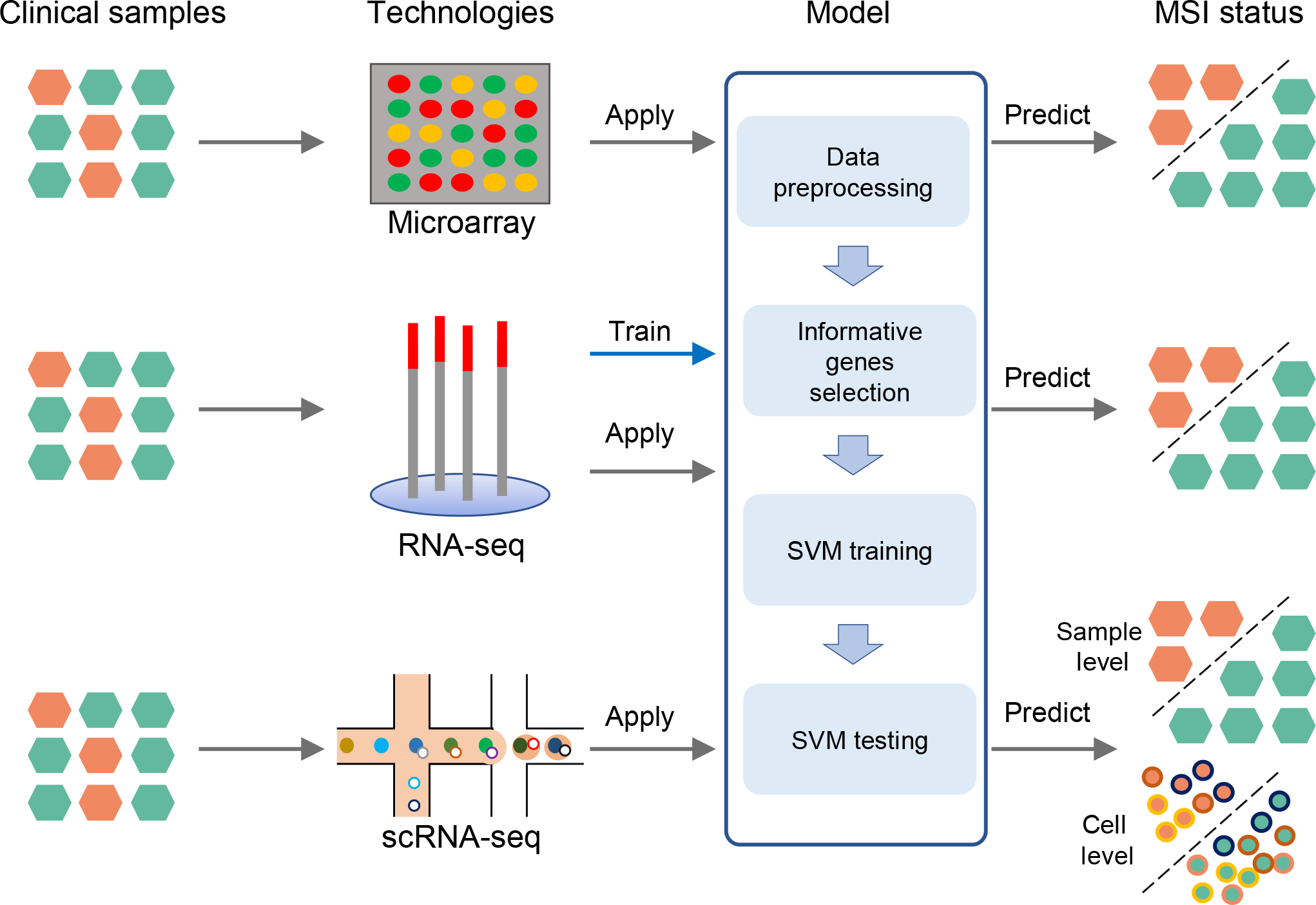
Workflow of MSIsensor-RNA. MSIsensor-RNA includes four modules: data preprocessing, informative gene selection, SVM model training, and testing. MSIsensor-RNA selects informative genes and trains SVM model by RNA-seq samples from TCGA. MSI scores are predicted by the trained model for microarray, RNA-seq, and scRNA-seq samples. SVM, support vector machine; scRNA-seq, single-cell RNA-seq, MSI, microsatellite instability.

### Data preprocessing

In MSIsensor-RNA, we accept microarray expression value, fragments per kilobase million (FPKM), transcripts per million (TPM), and RNA-Seq by expectation-maximization (RESM) read count as input. All expression matrix values were increased by 1 and then subjected to a log^2^ transformation. Then, for each sample or cell, the expression values were normalized to follow a Gaussian distribution with a mean of 0 and a standard deviation of 1. For the scRNA-seq sample, to obtain accurate MSI status, we only included high-quality cells with at least 20% of genes detected for MSI detection. If the number of high-quality cells was less than 20, all cells were sorted by the ratio of detected genes in descending order, and the top 20 cells would be utilized for MSI detection. To solve the dropout problem of scRNA-seq, zero value was imputed by the average of the gene expression values in the given sample.

### Informative gene selection

We selected informative genes for MSI classification in terms of stability, discrimination, and generalization, utilizing a dataset of 1428 TCGA samples with z-score-transformed gene expression values. Firstly, we removed ribosomal genes, mitochondrial genes, and genes with low FPKMs in TCGA dataset. Secondly, we selected genes with discriminative gene expression signatures between MSI samples and microsatellite stable (MSS) samples. We performed rank-sum test for expression values between MSI samples and MSS samples for each gene, and only genes with *P* < 0.01 (two-sided Wilcoxon rank-sum test) were included for the following analysis. Furthermore, we computed the log^2^ fold change of the *i*th gene as in Equation (1).

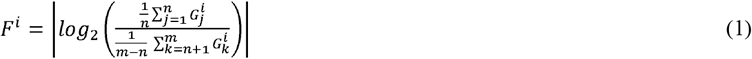

where *m* is the sample number for informative gene selection, *n* is the MSI sample number, and 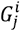 is the expression value of the *i*th gene for the *j*th sample. We only retained genes with a log^2^ fold change > 0.5 for the next step. We next calculated the AUC for each gene, and only genes with AUCs > 0.65 were regarded as candidate informative genes. To evaluate the generalization ability of the candidate informative genes, we employed both support vector machine (SVM) and random forest algorithms for each gene and calculated their respective 10-fold cross-validation scores. We then obtained two sorted gene lists based on the cross-validation results. Finally, we included the genes in the top 25% of both gene lists in the final informative gene set (Figure S2).

### Machine learning model training and testing

We build a SVM model to classify the MSI status for CRC, STAD, and UCEC in TCGA dataset. Firstly, we utilized SOMTE [27] to correct the imbalance between MSI and MSS in each cancer type by amplifying the MSI samples. Then, we utilized the expression values from the correct data as input to train the SVM model for MSI classification. To evaluate the performance of MSIsensor-RNA, we tested the trained model with 1848 independent samples from multiple platforms, including 247 RNA-seq, 1468 microarray, and 133 scRNA-seq samples. For a scRNA-seq sample, we calculated the MSI score with the SVM model for each high-quality cell. Then the average cell MSI score was used to evaluate the MSI status of a scRNA-seq sample.

### PreMSIm running

To compare the performance of MSIsensor-RNA with the only standalone software, PreMSIm, we also applied the data of microarray, RNA-seq, and scRNA-seq from 1848 independent samples to PreMSIm. For microarray and RNA-seq samples, we tested PreMSIm with two modes: PreMSIm-all and PreMSIm-split. In PreMSIm-all mode, we integrated all input samples from different databases and executed PreMSIm normalized and predicted modules. Conversely, the PreMSIm-split mode involved running PreMSIm separately for each database.

### Performance comparison of MSIsensor-RNA and PreMSIm

In MSIsensor-RNA, the predicted MSI probability by the SVM model was used to score the MSI status. The probabilities were further transformed into MSI status by the Youden index [28]. We first compared the MSIsensor-RNA score between MSI and MSS samples to test the performance of MSIsensor-RNA in multiple platforms using a rank-sum test. To further evaluate the performance of both two MSI detection methods, we calculated AUC, accuracy, F-score, precision, sensitivity, and specificity of MSIsensor-RNA and PreMSIm across different sequencing platforms.

### Robustness testing of MSIsensor-RNA and PreMSIm

To test the performance of MSIsensor-RNA and PreMSIm at different normalization methods, we tested these two methods with FPKM, TPM, and RESM read count formats of TCGA samples and calculated the AUC, F1-score, accuracy, precision, sensitivity, and specificity for each normalization method. To overcome the bias of different normalization methods and sequencing technologies, we normalized the input data of each sample to a Gaussian distribution with a mean of 0 and a standard deviation of 1. However, in PreMSIm, the normalization process was performed by genes, which means the normalized input data of a sample would be influenced by other samples in the bulk. Here, we tested the PreMSIm in two ways. Firstly, we input TCGA samples by three cancer types and calculated the performance of the predicted MSI status. Secondly, we input all TCGA samples together to evaluate their performance. We further compared the MSI result and performance of these two methods and found that the performance of PreMSIm was affected by the way input was provided.

## Results and discussion

The workflow of MSIsensor-RNA includes four modules (Figure 1 and Figure S1). First, we preprocess the expression value of microarray, bulk RNA-seq, and scRNA-seq data. Then, we select a set of informative genes for MSI detection. Next, we train a SVM model to estimate MSI scores using the gene expression values of the selected informative genes. Finally, we apply the trained model to predict MSI scores for either one clinical sample or a single cell (Table S1). For a given scRNA-seq sample, we also developed a model to report the MSI status of this sample by integrating MSI scores of cells within.

We first preprocessed all gene expression data and then applied the gene selection module to 1428 samples (the Z-score transferred gene expression values) from three MSI-popular cancer types (CRC, STAD, and UCEC) in TCGA dataset. Finally, we obtained 109 informative genes for MSI classification. We also performed this step for each type of CRC, STAD, and UCEC, yielding 397, 206, and 86 informative genes, respectively (Figures S3 and S4; Tables S2–S5). We found that only eight informative genes were detected in all three cancer types. Among these, we found that *MLH1* was the most important informative gene for MSI detection, as confirmed by previous reports [15,18,19] (Figure S5).

To assess the performance of MSIsensor-RNA in bulk sample data, we first trained tumor-specific models for CRC, STAD, and UCEC, as well as a model for all three MSI-popular cancer types in the TCGA dataset. Then, we compared the two kinds of models (tumor-specific and MSI-popular) with the standalone software, PreMSIm, in terms of the AUC, accuracy, sensitivity, and specificity in 1715 (1468 microarray and 247 bulk RNA-seq samples) independent samples. Notably, MSIsensor-RNA normalizes the expression value of informative genes for each sample independently, while PreMSIm must normalize each gene for multiple samples at the same time. Thus, we examined PreMSIm using both PreMSIm-all and PreMSIm-split modes.

For microarray data, we computed MSI status by MSIsensor-RNA and PreMSIm in 1468 samples from 12 GEO accessions. The result showed that MSIsensor-RNA predicted MSI with 0.952 AUC, while PreMSIm only performed 0.628 AUC in PreMSIm-split mode and 0.912 AUC in PreMSIm-all mode (**Figure 2A**; Figures S6 and S7; Tables S6 and S7). Meanwhile, MSIsensor-RNA achieved much higher sensitivities than PreMSIm-split and preMSIm-all (MSIsensor-RNA: 0.968, PreMSIm-split: 0.912, PreMSIm-all: 0.384), and performed comparable specificities with PreMSIm-split and preMSIm-all (MSIsensor-RNA: 0.843, PreMSIm-split: 0.912, PreMSIm-all: 0.873).

**Figure 2.**
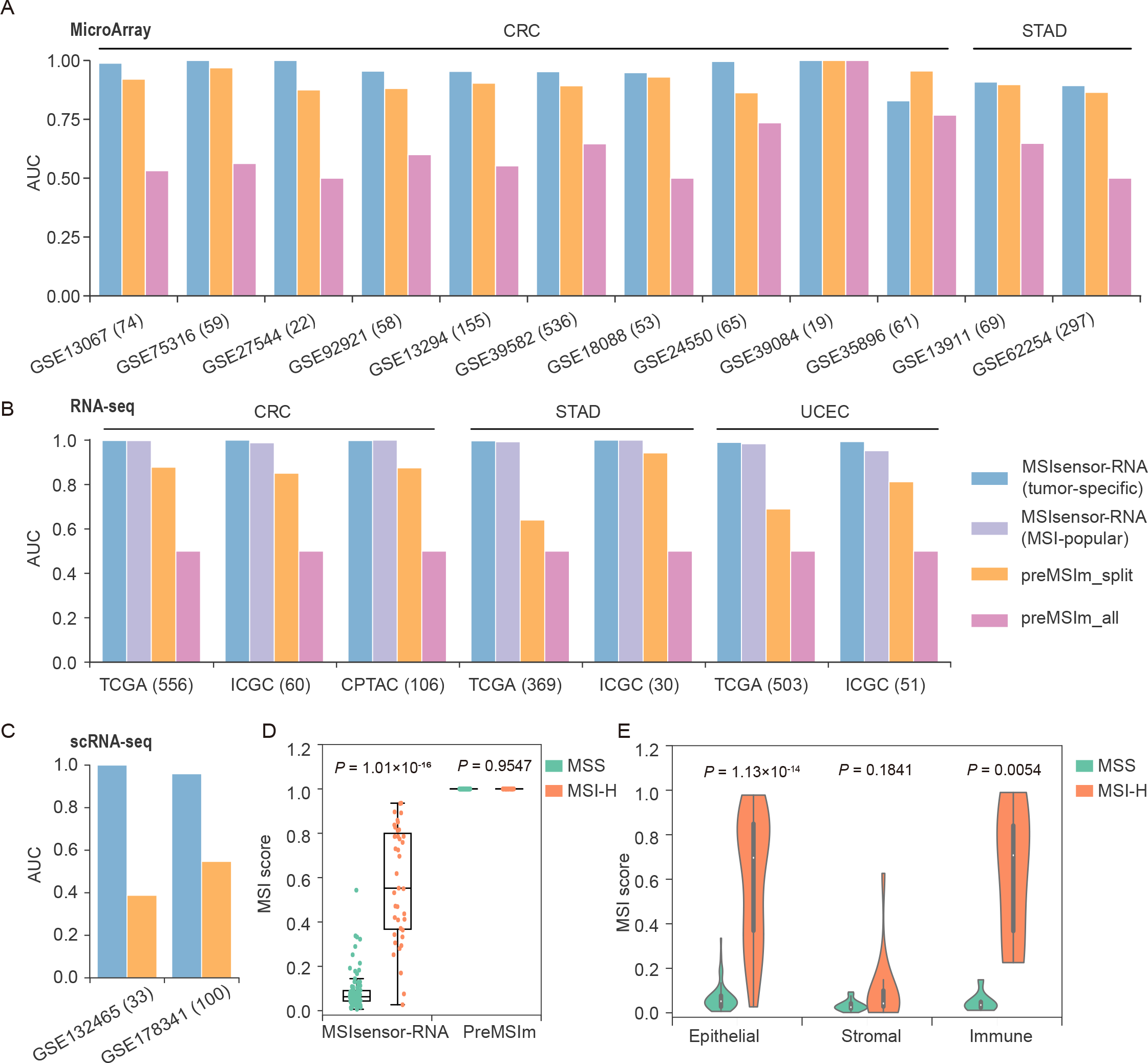
Performance of MSIsensor-pro A□C. UC of MSIsensor-RNA and PreMSIm in microarray (**A**), RNA-seq (**B**), and scRNA-seq (**C**) samples. In PreMSIm-all mode, all input samples from different databases are integrated for PreMSIm running. Conversely, the PreMSIm-split mode involves running PreMSIm separately for each database. **D**. Boxplot of MSIsensor-RNA score in scRNA-seq samples. **E**. Violin plot of MSIsensor-RNA score of different cell types in scRNA-seq samples. Epithelial, stromal, and immune cell types are defined in Pelka et al. [25]. *P* values were obtained through a two-sided rank-sum test. Tumor-specific, MSI results with the tumor-specific model; MSI-popular, MSI results with three MSI-popular cancer types; AUC, the area under the receiver operating characteristic curve; CRC, colorectal cancer; STAD, stomach adenocarcinoma; UCEC, uterine corpus endometrial carcinoma.

To evaluate the performance using bulk RNA-seq data, we compared MSIsensor-RNA and two modes of PreMSIm on 247 independent samples from ICGC and CPTAC. We noticed that MSIsensor-RNA achieved 0.997 AUC in the tumor-specific model and 0.985 AUC in the MSI-popular model, which were significantly greater than PreMSIm-all (0.5) and PreMSIm-split (0.870) (Figure 2B; Figures S8 and S9; Tables S8 and S9). In addition, MSIsensor-RNA performed much better than PreMSIm for both sensitivity (MSIsensor-RNA with tumor-specific model: 0.951, MSIsensor-RNA with MSI-popular model: 0.973, PreMSIm-split: 0.834, PreMSIm-all: 0.25) and specificity (MSIsensor-RNA with tumor-specific mode: 1, MSIsensor-RNA with MSI-popular model: 0.923, PreMSIm-split: 0.906, PreMSIm-all: 0.75). To further investigate the robustness of MSIsensor-RNA for different input data types, we evaluated the performance of MSIsensor-RNA and PreMSIm with FPKM, read count, and TPM normalized samples in TCGA as input. We found that MSIsensor-RNA achieved 0.982 ± 0.040 AUC, indicating its robustness regardless of the measurement methods of gene expression (Table S10).

To assess the performance of MSIsensor-RNA and PreMSIm in scRNA-seq samples, we applied the trained model from TCGA dataset to 23,902 high-quality cells from 133 samples to obtain sample-specific MSI status and compared it to the ratio of cells labeled as MSI by PreMSIm. The result indicated that MSIsensor-RNA detected MSI for scRNA-seq samples with 0.958 AUC, 0.9231 sensitivity, and 0.9362 specificity, while PreMSIm had 0.4969 AUC, 1 sensitivity, and 0.0319 specificity (Figure 2C; Figure S10; Tables S11 and S12). The sample level MSI scores based on scRNA-seq were significantly different between MSI and MSS samples by MSIsensor-RNA (two-sided rank-sum test, *P* = 1.01×10^−16^), while no significant difference was detected for PreMSIm (two-sided rank-sum test, *P* = 0.9547) (Figure 2D). Having established the effectiveness of MSIsensor-RNA on scRNA-seq samples, we investigated cell-level MSI. We computed the MSI scores of 21,438 high-quality cells from 100 samples (GSE178341) and found MSI score was correlated with cell type. For example, MSI scores of epithelial and immune cells in MSI samples were greater than those in MSS samples, while no significant difference was detected between MSI and MSS for stromal cells (Figure 2E; Figure S11; Table S13). This indicated the potential of MSIsensor-RNA to assess MSI at the single-cell level, providing a novel measurement for the investigation of tumorigenic processes.

Microsatellite instability is important for the prognosis assessment of both 5-FU chemotherapy [4] and immunotherapy [5]. In addition to gold-standard experimental methods [6], MSI status is also evaluated according to genomic sequencing data [7-14], gene expression data [15,18,19], methylation data [17], as well as hematoxylin and eosin-stained slides [20,21]. Compared to variants in microsatellite regions, gene expression values are more directly reflective of the features of MSI and easier to obtain. In this study, we have developed a robust method, MSIsensor-RNA, for MSI detection with gene expression data. MSIsensor-RNA provides informative gene selection, model training, and MSI detection modules and is able to process data from multiple platforms, including microarray, RNA-seq, and single-cell RNA-seq. Compared to the standalone method PreMSIm, MSIsensor-RNA also provides modules for informative gene selection and model training, enabling users to apply it across different cancer types. MSIsensor-RNA also improves the normalization method of the data, yielding a more robust result than PreMSIm (Figure 2). In addition, MSIsensor-RNA facilitates the evaluation of MSI status at the single cell level, which will be critical to better understanding the mechanism of MSI in cancer immunotherapy in the future.

In most MSI detection methods, such as MSIsensor [10] and MSIsensor-pro [11], MSI is quantified according to genetic mutations at microsatellite sites, the consequence of MSI, rather than the deficiency of the MMR system, the direct cause of MSI. In this study, a set of MSI-associated genes was identified, and their expression values were used for MSI evaluation. We found that *MLH1* is the most important gene in all tested cancer types. In addition, unexpected expression of *MLH1* is commonly seen in Lynch syndrome [29]. Thus, we test the performance of MSIsensor-RNA for samples with abnormal *MLH1* expression. We trained a model based on all informative genes and tested it using samples with simulated abnormal *MLH1* gene expression (Table S14). We found that the model achieved 0.974 and 0.972 AUCs when we set the *MLH1* expression value as the maximum and minimum of all gene expression values, respectively. Furthermore, when *MLH1* was excluded from the informative gene set, MSIsensor-RNA also achieved a 0.977 AUC, indicating the robustness of MSIsensor-RNA for MSI detection.

We demonstrated that MSIsensor-RNA achieved higher performance than other methods based on gene expression and comparable performance compared to DNA-based methods (Table S15). In our study design, MSIsensor-RNA detects MSI according to the gene expression signature of genes on MSI-associated pathways, while MSIsensor evaluates MSI by computing the ratio of somatic microsatellite mutations. Although MSIsensor achieved slightly higher performance than MSIsensor-RNA, it cannot replace the applications of MSIsensor-RNA in gene expression data. Currently, MSIsensor-RNA reports favorable performance in all three MSI-popular cancers, including colorectal cancer, stomach adenocarcinoma, and uterine corpus endometrial carcinoma. The MSI features vary across different cancer types. Thus, the model obtained low performance when the testing samples were inconsistent with training samples in cancer types (Tables S16□S18). Therefore, the performance of MSIsensor-RNA in other cancer types needs further validation in the future. Another characteristic of MSIsensor-RNA is that its performance is not stable when trained with only a small number of positive samples. For example, in gastric cancer, the MSI-H training samples in stages I and IV are much fewer compared to other stages, leading to lower performance in these two stages (Tables S19 and S20).

## Conclusions

MSIsensor-RNA is a cross-platform, efficient, and robust method for MSI status determination from both bulk and single-cell gene expression data. We demonstrated the effectiveness and robustness of MSIsensor-RNA across different platforms, hinting at its potential in clinical research. Moreover, MSIsensor-RNA enables single-cell level MSI evaluation, providing a new tool to discover the role of MSI in the tumorigenic process and to monitor cell-level dynamic changes during immunotherapy.

## Code availability

MSIsensor-RNA is available at https://ngdc.cncb.ac.cn/biocode/tools/7385 and https://github.com/xjtu-omics/msisensor-rna.

## CRediT authorship statement

**Peng Jia:** Methodology, Software, Validation, Formal analysis, Investigation, Visualization, Writing - Original Draft, Writing - Review & Editing. **Xuanhao Yang:** Methodology, Software, Formal analysis, Investigation, Data Curation, Writing - Original Draft. **Xiaofei Yang:** Methodology, Funding acquisition. **Tingjie Wang:** Methodology. **Yu Xu:** Methodology. **Kai Ye:** Methodology, Resources, Conceptualization, Supervision, Project administration, Funding acquisition.

## Acknowledgments

KY and XY are supported by the National Key R&D Program of China (Grant No. 2022YFC3400300), the National Natural Science Foundation of China (Grant Nos. 32125009, 32070663, and 62172325), the Fundamental Research Funds for the Central Universities (Grant No. xzy012020012), and the Natural Science Basic Research Program of Shaanxi (Grant No. 2021GXLH-Z-098), China. We thank everyone in the Ye lab at Xi’an Jiaotong University for their helpful comments and discussion. We also thank the producers of the data we used in this study.

## Supplementary material

**Figure S1 Workflow of MSIsensor-RNA**. The diagram illustrates the detailed steps of training and testing for MSIsensor-pro. WES, whole exome sequencing.

**Figure S2 Workflow of informative gene selection**. MSIsensor-RNA selects informative genes based on criteria such as stability, discrimination, and generalization. Initially, mitochondrial genes and ribosomal genes are excluded from consideration. Subsequently, genes with expression values that do not exhibit significant differences between MSI and MSS samples (as determined by a two-sided Wilcoxon rank-sum test) are filtered out. Finally, genes with expression values that demonstrate a high generalization score for MSI detection are retained. MSI, microsatellite instability; MSS, microsatellite stable; AUC, the area under the receiver operating characteristic curve; SVM, support vector machine.

**Figure S3 Venn plot showing the intersection of the informative genes across different cancer types**. The pink circle represents the informative genes generated from samples of three MSI-popular cancer types including CRC, STAD, and UCEC. CRC, colorectal cancer; STAD, stomach adenocarcinoma; UCEC, uterine corpus endometrial carcinoma.

**Figure S4 Upset plot showing the intersection of the informative genes across this study and the other three publications**. All four studies utilized *MLH1* as an informative gene.

**Figure S5 Violin plot of expression values of mismatch repair genes in MSI and MSS samples**. A two-sided Wilcoxon rank-sum test is implemented to compare MSI scores between MSI and MSS samples. FPKM, fragments per kilobase million; ns, not significant; *, *P* < 0.0001.

**Figure S6 Violin plot of MSI score by MSIsensor-RNA in microarray data**. The violin plot shows that MSI scores by MSIsensor-RNA in MSI are significantly greater than those in MSS samples. A two-sided Wilcoxon rank-sum test is implemented to compare MSI scores between MSI and MSS samples. ns, not significant; *, *P* < 0.05; **, P < 0.01; ***, P < 0.001; ****, *P* < 0.0001.

**Figure S7 Performance of MSIsensor-RNA, PreMSIm-all, and PreMSIm-split for microarray data**. The bar plot shows that MSIsensor-RNA outperforms PreMSIm-all and PreMSIm-split across AUC, accuracy, sensitivity, and specificity. In PreMSIm-all mode, all input samples from different databases are integrated for PreMSIm running. Conversely, the PreMSIm-split mode involves running PreMSIm separately for each database.

**Figure S8 Violin plot of MSI score by MSIsensor-RNA in bulk RNA-seq data**. The violin plot shows that MSI scores by MSIsensor-RNA in MSI are significantly greater than those in MSS samples. A two-sided Wilcoxon rank-sum test is implemented to compare MSI scores between MSI and MSS samples.

**Figure S9 Performance of MSIsensor-RNA, PreMSIm-all, and PreMSIm-split for bulk RNA-seq data**. The bar plot shows that MSIsensor-RNA outperforms PreMSIm-all and PreMSIm-split across AUC, accuracy, sensitivity, and specificity in all three databases. PreMSIm performs nearly 100% sensitivities but no more than 50% specificities in TCGA and ICGC. Meanwhile, PreMSIm predicts MSI status with 100% specificities but 0.7 sensitivities in PreMSIm-split and 0 in PreMSIm-all.

**Figure 10 Performance of MSIsensor-RNA, PreMSIm-all, and PreMSIm-split for scRNA-seq data**. The bar plot shows that MSIsensor-RNA outperforms PreMSIm-all and PreMSIm-split across AUC, accuracy, sensitivity, and specificity in the two databases.

**Figure S11 Scatter plot shows the distribution of MSI score and cell ratio of differential cell types in GSE178341**. The result shows that epithelial cell is a large proportion of tumor samples, while stromal cell and immune cell are only less than 25%. The MSI scores of epithelial cells and immune cells in MSI samples are significantly greater than those in MSS samples, but MSI scores in stromal cells have no significant difference between MSI and MSS samples.

**Table S1 Overview of samples in this study**

**Table S2 Details of informative genes in colorectal cancer**.

**Table S3 Details of informative genes in stomach adenocarcinoma**.

**Table S4 Details of informative genes in uterine corpus endometrial carcinoma**.

**Table S5 Details of informative genes in three MSI-popular cancers**.

**Table S6 MSI results of microarray samples by MSIsensor-RNA and PreMSIm**.

**Table S7 MSI detection performance of MSIsensor-RNA and PreMSIm in microarray samples**.

**Table S8 MSI results of RNA-seq samples by MSIsensor-RNA and PreMSIm. Table S9 MSI detection performance of MSIsensor-RNA and PreMSIm in RNA-seq samples**.

**Table S10 MSI detection performance of MSIsensor-RNA and PreMSIm in different normalized samples**.

**Table S11 MSI results of scRNA-seq samples by MSIsensor-RNA**.

**Table S12 MSI detection performance of MSIsensor-RNA and preMSIm in scRNA-seq samples**.

**Table S13 MSI results of scRNA-seq cells by MSIsensor-RNA**.

**Table S14 Performance of MSIsensor-RNA with abnormal *MLH1* expression values**.

**Table S15 Performance of MSIsensor-RNA and MSIsensor in TCGA dataset. Table S16 AUC of MSIsensor-RNA with inconsistent training and testing samples**.

**Table S17 Performance of train models for cancer with low-frequency MSI. Table S18 Performance of MSIsensor-RNA for cancer with low-frequency MSI by 5-fold cross-validation**.

**Table S19 Performance of MSIsensor-RNA across different cancer stages. Table S20 Summary of MSI-H training samples for STAD in Table S19**.

## References

[1] Yamamoto H, Imai K. Microsatellite instability: an update. Arch Toxicol 2015;89:899□921.

[2] Hause RJ, Pritchard CC, Shendure J, Salipante SJ. Classification and characterization of microsatellite instability across 18 cancer types. Nat Med 2016;22:1342□50.

[3] Rizvi NA, Hellmann MD, Snyder A, Kvistborg P, Makarov V, Havel JJ, et al. Cancer immunology. Mutational landscape determines sensitivity to PD-1 blockade in non-small cell lung cancer. Science 2015;348:124□8.

[4] Ribic CM, Sargent DJ, Moore MJ, Thibodeau SN, French AJ, Goldberg RM, et al. Tumor microsatellite-instability status as a predictor of benefit from fluorouracil-based adjuvant chemotherapy for colon cancer. N Engl J Med 2003;349:247□57.

[5] Le DT, Uram JN, Wang H, Bartlett BR, Kemberling H, Eyring AD, et al. PD-1 Blockade in tumors with mismatch-repair deficiency. N Engl J Med 2015;372:2509□20.

[6] Boland CR, Thibodeau SN, Hamilton SR, Sidransky D, Eshleman JR, Burt RW, et al. A national cancer institute workshop on microsatellite instability for cancer detection and familial predisposition: development of international criteria for the determination of microsatellite instability in colorectal cancer. Cancer Res 1998;58:5248□57.

[7] Kautto EA, Bonneville R, Miya J, Yu L, Krook MA, Reeser JW, et al. Performance evaluation for rapid detection of pan-cancer microsatellite instability with MANTIS. Oncotarget 2017;8:7452□63.

[8] Niu B, Ye K, Zhang Q, Lu C, Xie M, McLellan MD, et al. MSIsensor: microsatellite instability detection using paired tumor-normal sequence data. Bioinformatics 2014;30:1015□6.

[9] Salipante SJ, Scroggins SM, Hampel HL, Turner EH, Pritchard CC. Microsatellite instability detection by next generation sequencing. Clin Chem 2014;60:1192□9.

[10] Jia P, Yang X, Guo L, Liu B, Lin J, Liang H, et al. MSIsensor-pro: fast, accurate, and matched-normal-sample-free detection of microsatellite instability. Genomics Proteomics Bioinformatics 2020;18:65□71.

[11] Hempelmann JA, Scroggins SM, Pritchard CC, Salipante SJ. MSIplus for integrated colorectal cancer molecular testing by next-generation sequencing. J Mol Diagn 2015;17:705□14.

[12] Maruvka YE, Mouw KW, Karlic R, Parasuraman P, Kamburov A, Polak P, et al. Analysis of somatic microsatellite indels identifies driver events in human tumors. Nat Biotechnol 2017;35:951□9.

[13] Huang MN, McPherson JR, Cutcutache I, Teh BT, Tan P, Rozen SG. MSIseq: software for assessing microsatellite instability from catalogs of somatic mutations. Sci Rep 2015;5:13321.

[14] Ratovomanana T, Cohen R, Svrcek M, Renaud F, Cervera P, Siret A, et al. Performance of next-generation sequencing for the detection of microsatellite instability in colorectal cancer with deficient DNA mismatch repair. Gastroenterology 2021;161:814□26.

[15] Pacinkova A, Popovici V. Cross-platform data analysis reveals a generic gene expression signature for microsatellite instability in colorectal cancer. Biomed Res Int 2019;2019:6763596.

[16] Middha S, Zhang L, Nafa K, Jayakumaran G, Wong D, Kim HR, et al. Reliable pan-cancer microsatellite instability assessment by using targeted next-generation sequencing data. JCO Precis Oncol 2017;2017:1□17.

[17] Foltz SM, Liang WW, Xie M, Ding L. MIRMMR: binary classification of microsatellite instability using methylation and mutations. Bioinformatics 2017;33:3799□801.

[18] Li L, Feng Q, Wang X. PreMSIm: An R package for predicting microsatellite instability from the expression profiling of a gene panel in cancer. Comput Struct Biotechnol J 2020;18:668□75.

[19] Danaher P, Warren S, Ong S, Elliott N, Cesano A, Ferree S. A gene expression assay for simultaneous measurement of microsatellite instability and anti-tumor immune activity. J Immunother Cancer 2019;7:15.

[20] Echle A, Grabsch HI, Quirke P, van den Brandt PA, West NP, Hutchins GGA, et al. Clinical-grade detection of microsatellite instability in colorectal tumors by deep learning. Gastroenterology 2020;159:1406□16 e11.

[21] Kather JN, Pearson AT, Halama N, Jager D, Krause J, Loosen SH, et al. Deep learning can predict microsatellite instability directly from histology in gastrointestinal cancer. Nat Med 2019;25:1054□6.

[22] Longo SK, Guo MG, Ji AL, Khavari PA. Integrating single-cell and spatial transcriptomics to elucidate intercellular tissue dynamics. Nat Rev Genet 2021;22:627□44.

[23] Fujimoto A, Fujita M, Hasegawa T, Wong JH, Maejima K, Oku-Sasaki A, et al. Comprehensive analysis of indels in whole-genome microsatellite regions and microsatellite instability across 21 cancer types. Genome Res 2020;30:334□346.

[24] Vasaikar S, Huang C, Wang X, Petyuk VA, Savage SR, Wen B, et al. Proteogenomic analysis of human colon cancer reveals new therapeutic opportunities. Cell 2019;177:1035□49 e19.

[25] Pelka K, Hofree M, Chen JH, Sarkizova S, Pirl JD, Jorgji V, et al. Spatially organized multicellular immune hubs in human colorectal cancer. Cell 2021;184:4734□52 e20.

[26] Lee HO, Hong Y, Etlioglu HE, Cho YB, Pomella V, Van den Bosch B, et al. Lineage-dependent gene expression programs influence the immune landscape of colorectal cancer. Nat Genet 2020;52:594□603.

[27] Chawla NV, Bowyer KW, Hall LO, Kegelmeyer WP. SMOTE: synthetic minority over-sampling technique. Journal of Artificial Intelligence Research 2002;16:321□57.

[28] Schisterman EF, Perkins NJ, Liu A, Bondell H. Optimal cut-point and its corresponding Youden Index to discriminate individuals using pooled blood samples. Epidemiology 2005;16:73□81.

[29] Chen W, Hampel H, Pearlman R, Jones D, Zhao W, Alsomali M, et al. Unexpected expression of mismatch repair protein is more commonly seen with pathogenic missense than with other mutations in Lynch syndrome. Hum Pathol 2020;103:34□41.

