## Supplementary material for "MSIsensor-RNA: Microsatellite Instability Detection for Bulk and Single-cell Gene Expression Data": Figure S1

### Slide 1
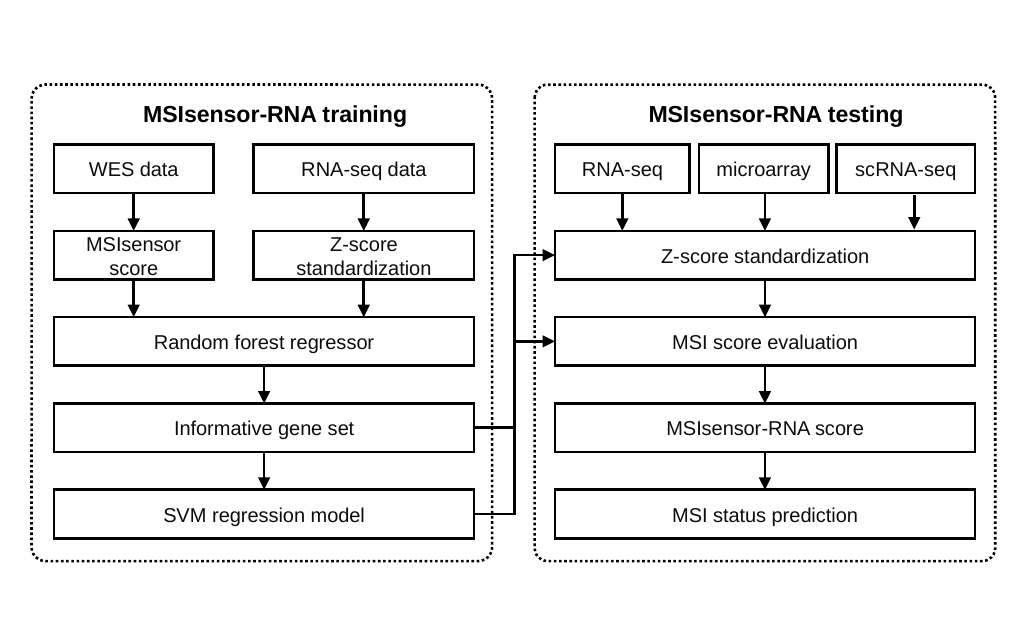

MSIsensor-RNA training
MSIsensor-RNA testing
WES data
RNA-seq data
RNA-seq
microarray
scRNA-seq
MSIsensor score
Z-score standardization
Z-score standardization
Random forest regressor
MSI score evaluation
Informative gene set
MSIsensor-RNA score
SVM regression model
MSI status prediction
