## Supplementary material for "MSIsensor-RNA: Microsatellite Instability Detection for Bulk and Single-cell Gene Expression Data": Figure S2

### Slide 1
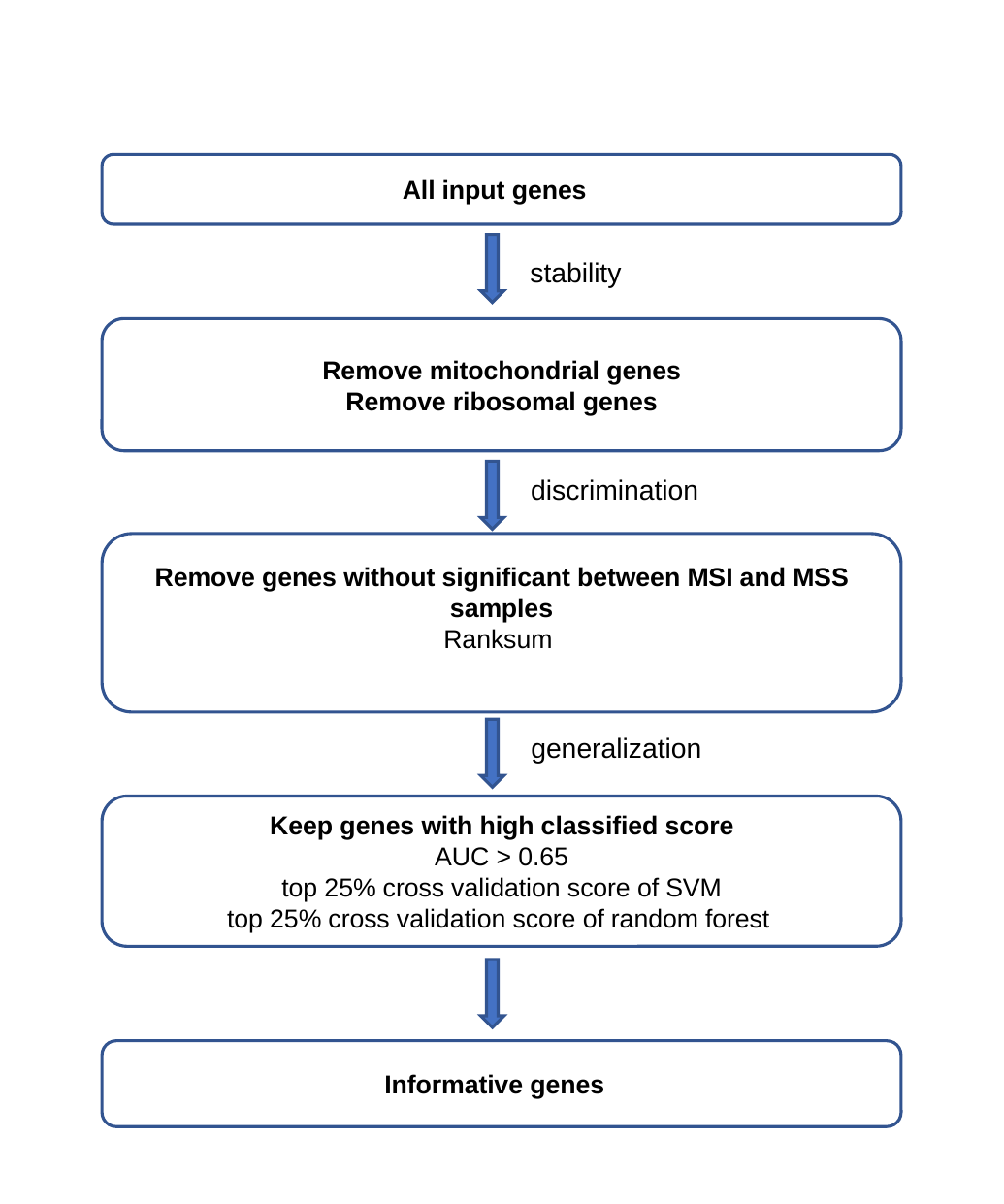

All input genes
stability
Remove mitochondrial genes
Remove ribosomal genes
discrimination
generalization
Keep genes with high classified score
AUC > 0.65
 top 25% cross validation score of SVM
top 25% cross validation score of random forest
Informative genes
