## Supplementary figures and images for "MSIsensor-RNA: Microsatellite Instability Detection for Bulk and Single-cell Gene Expression Data"

### Figure S3

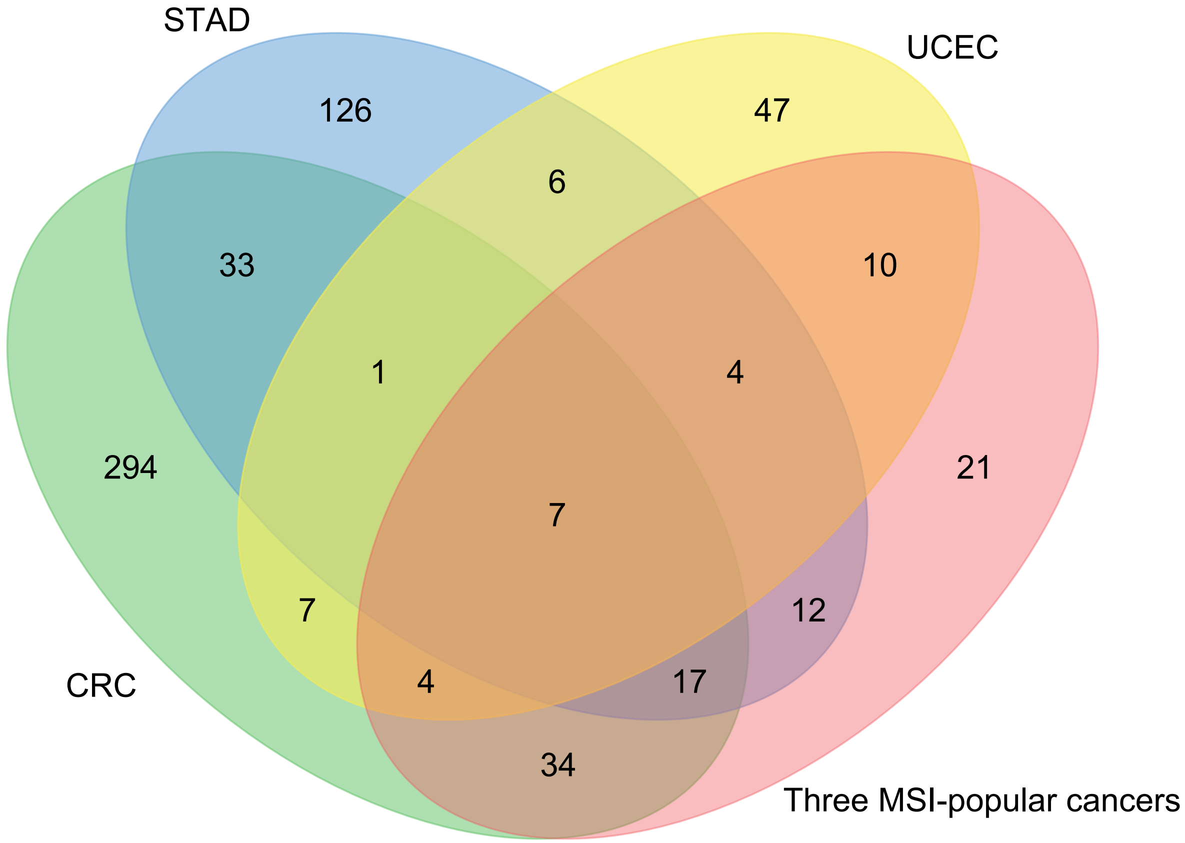

### Figure S4

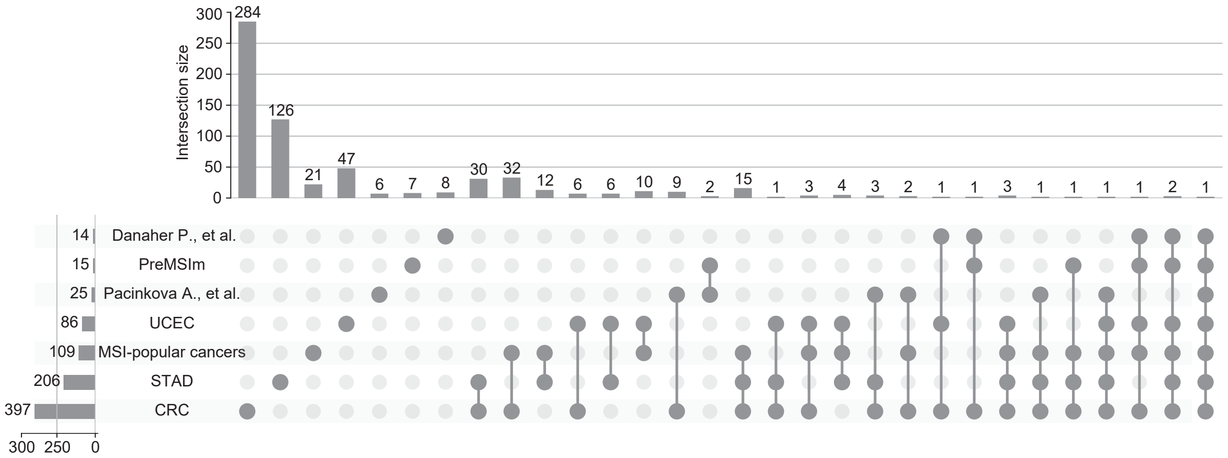

### Figure S5

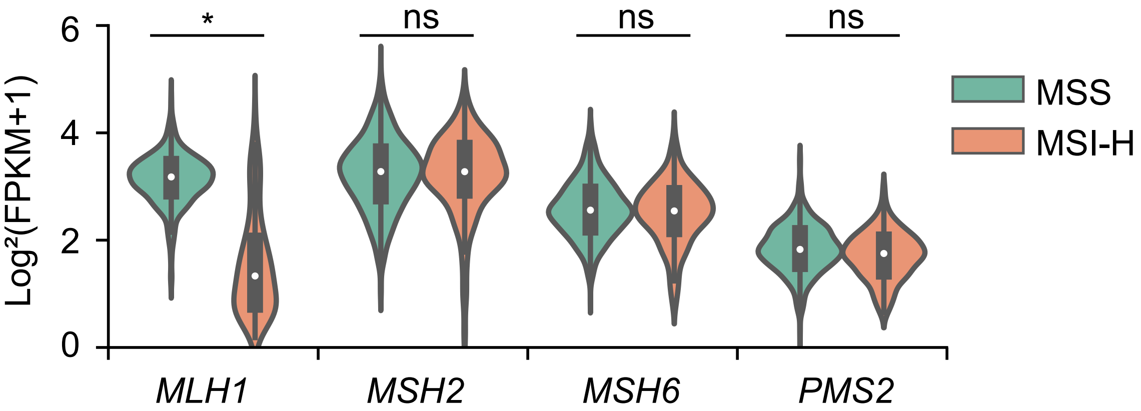

### Figure S6

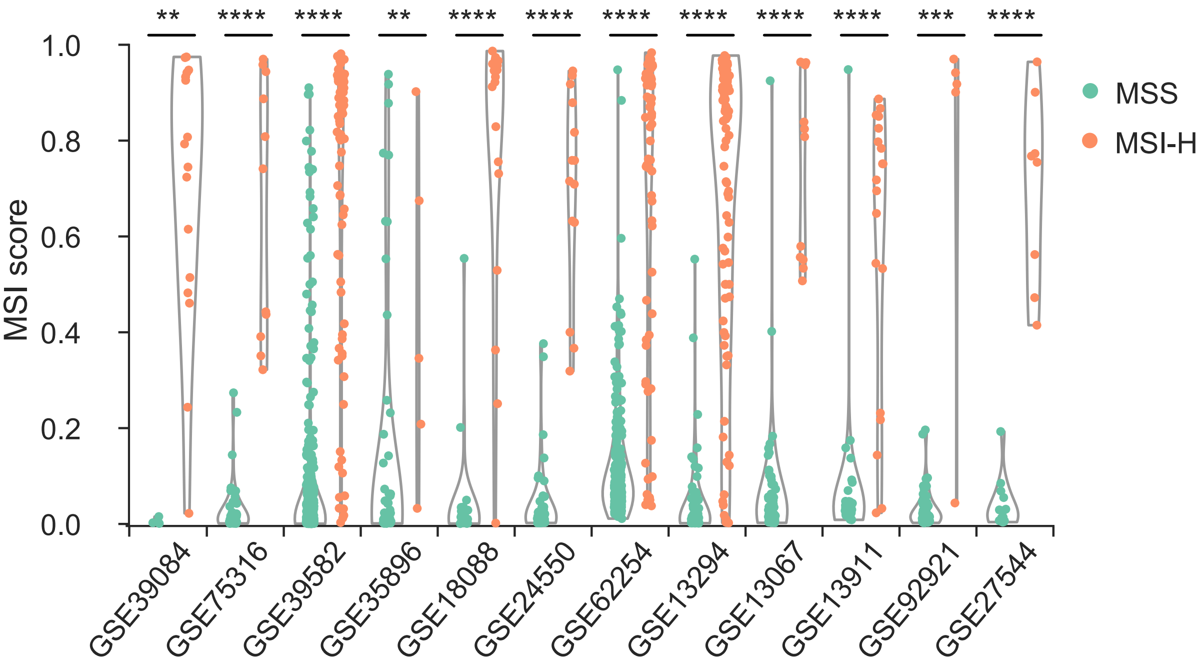

### Figure S7

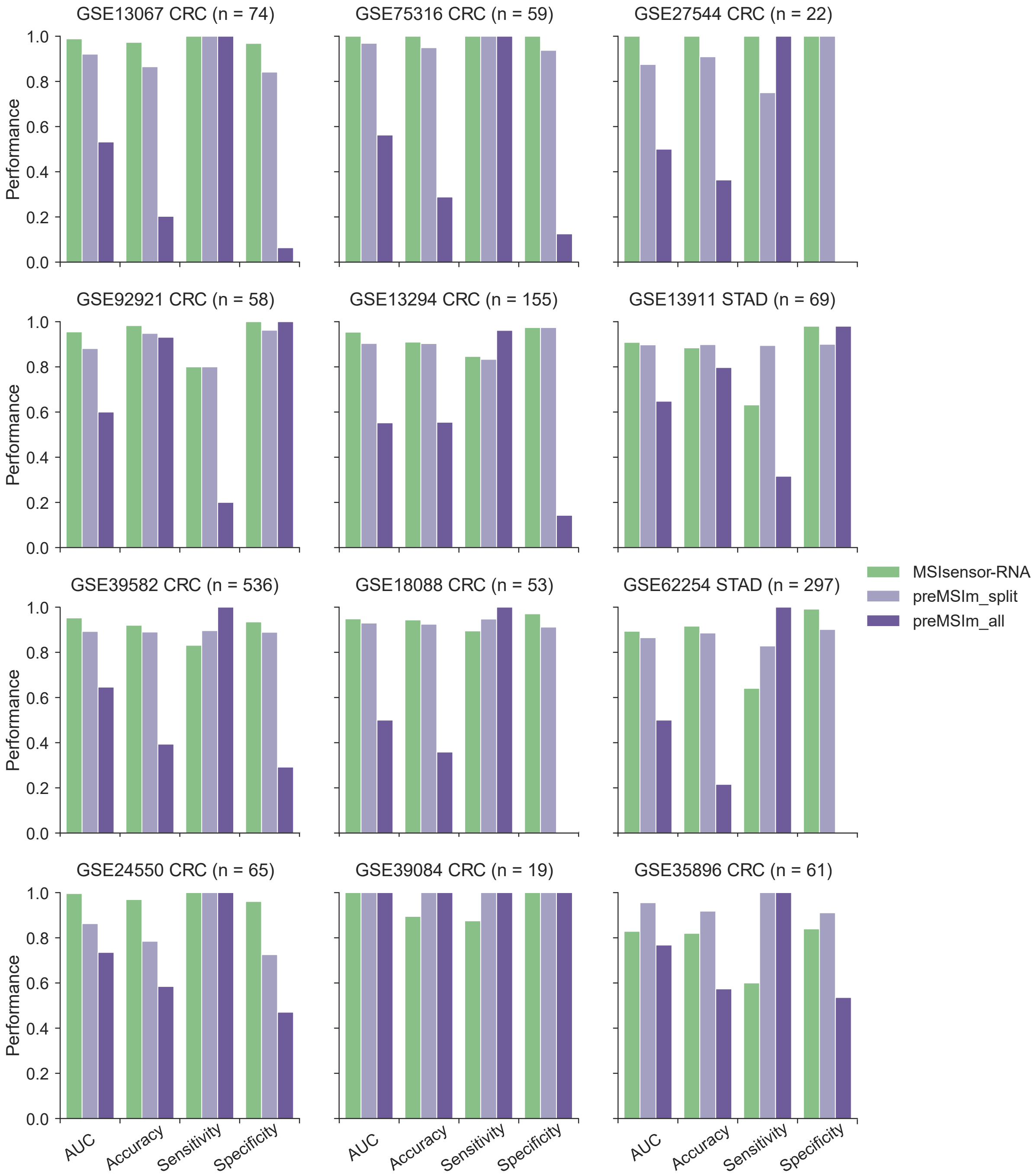

### Figure S8

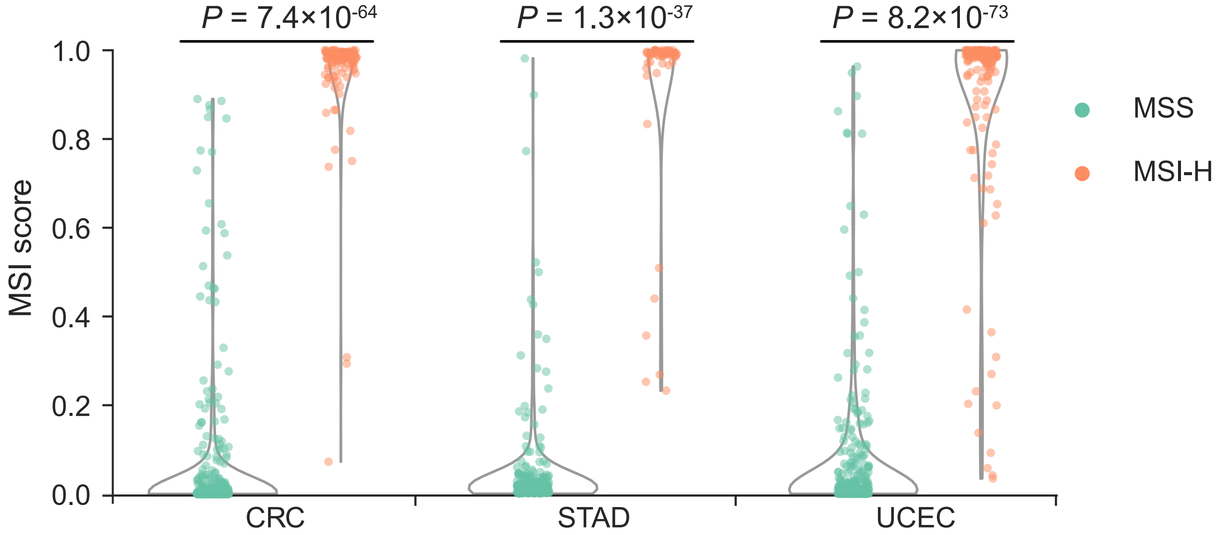

### Figure S9

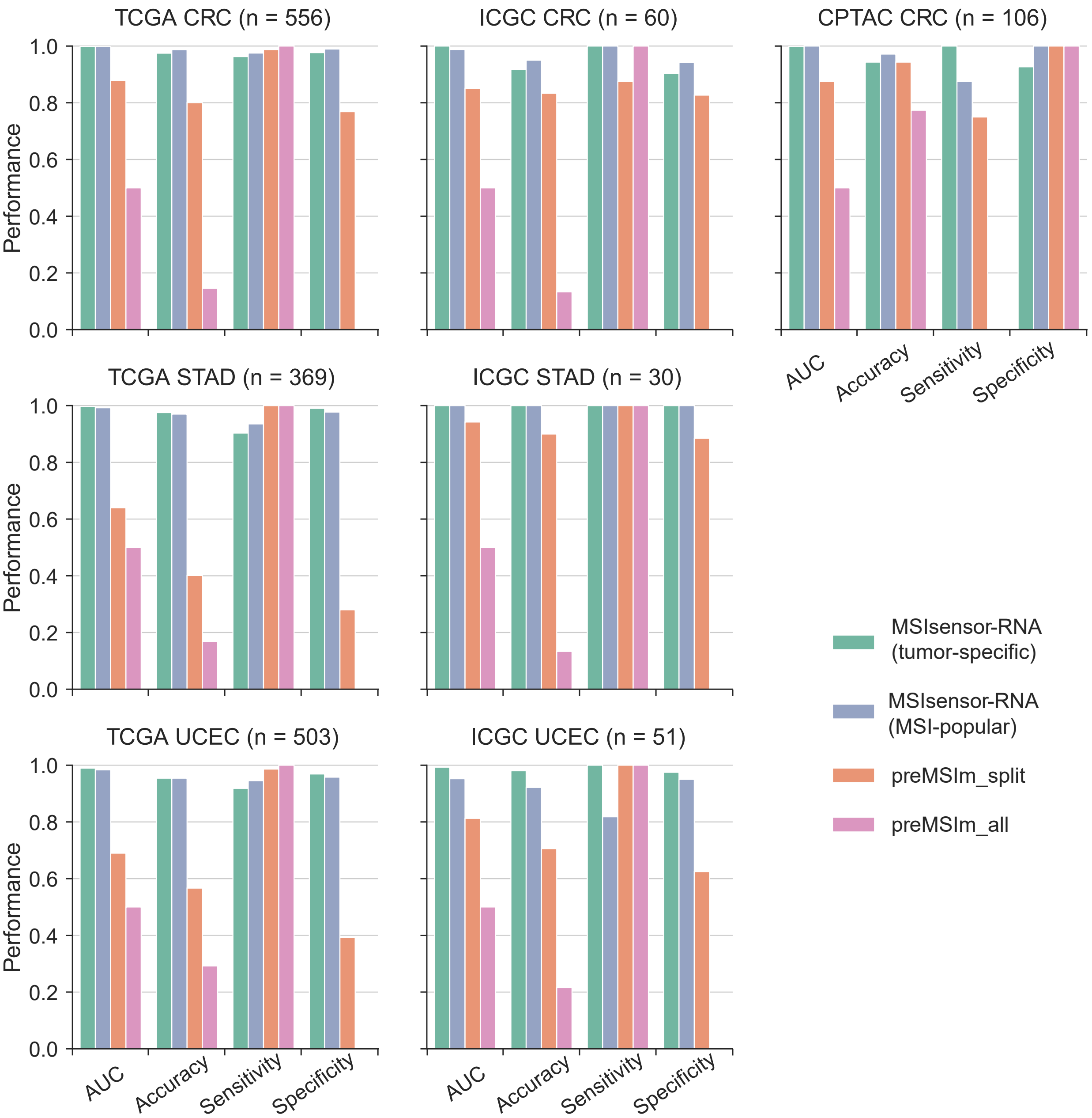

### Figure S10

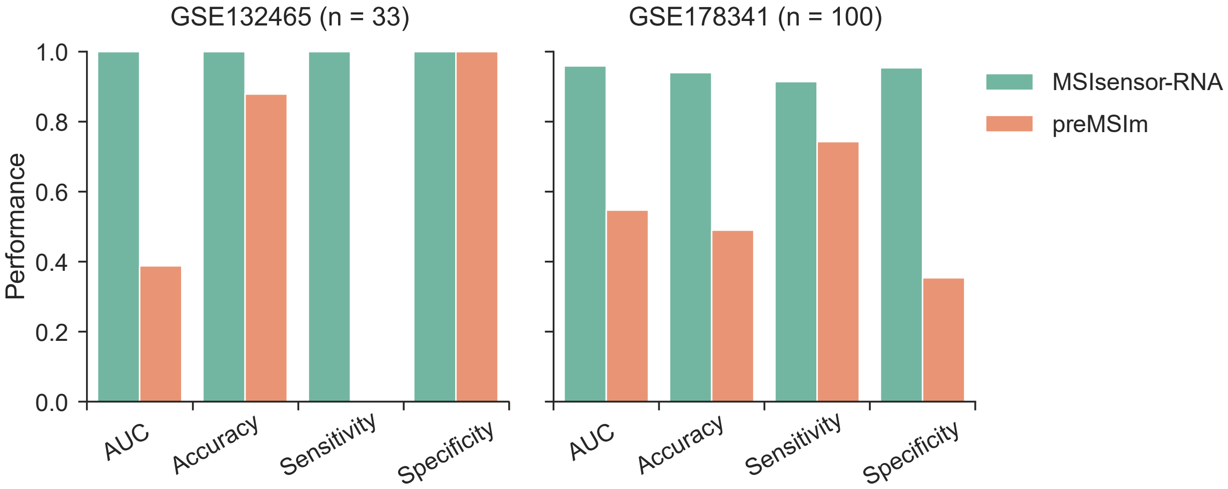

### Figure S11

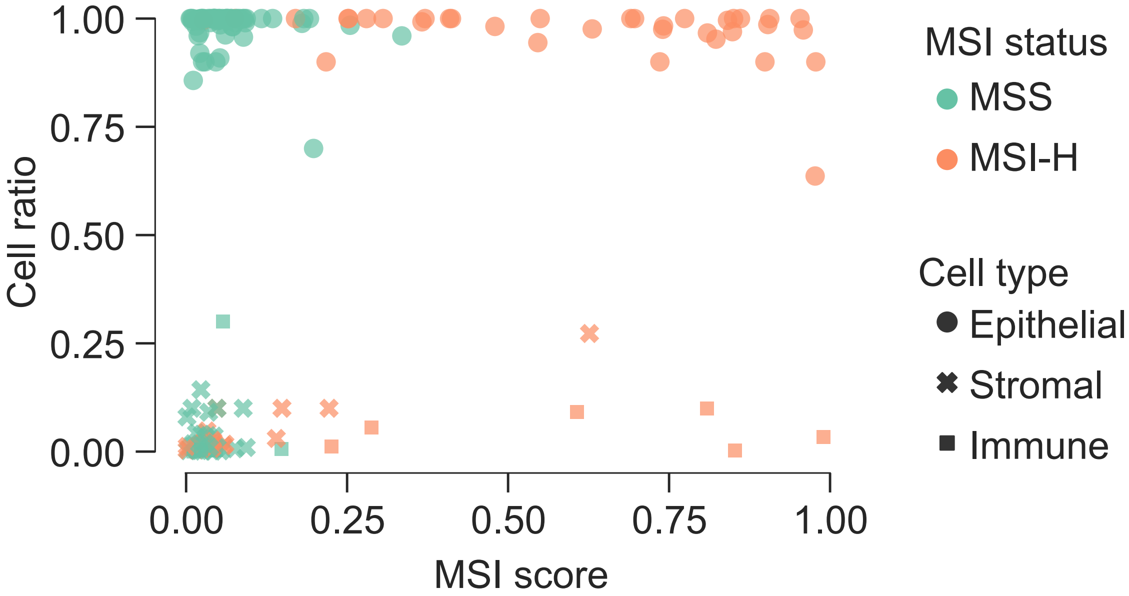
